# Comprehensive Analysis of the Expression and Prognosis for *PSMBs* in Clear Cell Renal Cell Carcinoma

**DOI:** 10.1101/2022.05.12.491733

**Authors:** Ning Jiang, Yulin Zhang, Zihan Zhao, Tianhang Li, Tianyao Liu, Hongqian Guo, Rong Yang

## Abstract

**Background:** Clear cell renal cell carcinoma (ccRCC) is one of the most common malignances with an ever-increasing incidence and high mortality. Although comprehensive therapy has made great progress in recent years, the prognosis of most ccRCC patients is still poor. The proteasome β subunits (*PSMBs*) are a group of genes composing the ubiquitin–proteasome pathway, which is the principal mechanism of protein catabolism in mammalian cytoplasm and nucleus and is essential for the regulation of almost all cellular processes. Growing evidence show s that the *PSMBs* play an important role in the occurrence and progression of various cancers. However, the expression and prognostic value of *PSMBs* in ccRCC have not been evaluated.

**Methods:** To solve this issue, ONCOMINE, UALCAN, Human Protein Atlas, cBioPortal, Metascape, GeneMANIA, STRING, DAVID 6.8, TIMER were used in this study.

**Results:** The transcriptional expression levels of *PSMB2/4/7/8/9/10* in ccRCC tissues were found to be significantly upregulated, and the mRNA expression levels of *PSMB1/2/3/4/8/9/10* were associated with advanced tumor stage and tumor grade. Besides, higher mRNA expression of *PSMB1/2/3/4/5/6/7/10* were associated with a shorter overall survival (OS). Furthermore, the protein expression level of *PSMB4/7/8/9/10* in ccRCC was higher than normal renal tissues. Significant relationships were also found between the expression of *PSMBs* and the infiltration of immune-related cells.

**Conclusions:** The present study implied that *PSMB4* and *PSMB10* might act as potential therapy targets and prognostic biomarkers in ccRCC, which may contribute to available knowledge, improve the treatment designs, and enhance the accuracy of prognosis for patients with ccRCC.

## 1 Introduction

Renal cell carcinoma (RCC) ranks as the 6th most commonly diagnosed cancer in men and 10th in women worldwide ^1^. Moreover, over 140000 RCC-related deaths occur yearly, which endows RCC with the 13th highest cancer-caused motility around the world ^2^. Clear cell renal cell carcinoma (ccRCC) is the most common malignances with an ever-increasing incidence and high mortality in RCC. Of note, because of the difficulty in distinguishing the primary lesion and the concealment of symptoms, nearly 1/5 patients have already developed distant metastases at first diagnosis^3^. Contemporary clinical therapies for ccRCC include surgery, chemotherapy, radiation therapy and immunotherapy ^4^. Despite the rapid progress made in the treatment of ccRCC, the survival rate of ccRCC patients is still far from expectations. High tumor heterogeneity and uncontrollability of tumor progression urge us to find more effective biomarkers to enhance individualized medical decision making and guide therapy selection more accurately.

Proteasome, functioning as a multisubunit cellular organelle, plays a major role in regulating protein degradation in the manner of a nonlysosomal threonine protease ^5^. Numerous studies suggested that protein degradation participates in various cellular biological processes, including cancer development ^6,7^. The proteasome is constituted by a 20S core complex along with a 19S regulatory complex. Two ring-like α-subunits and two β-subunits formed the 20Score component. Proteasome β subunit (PSMB) represents the family of the inner βrings with distinct proteolytic capabilities. *PSMB1-7* pointed to β1 to 7 subunits correspondingly. While *PSMB8-10* referred to β1i, β5i and β2i, which formed immunoproteasome in combination ^8^. To date, several *PSMBs* have been found to be involved in the tumor malignant biological behaviors. Wang, et al demonstrated that *PSMB5* served as oncogene in breast cancer and exerted immunosuppressive functions by inducing M1-to-M2 differentiation in the tumor micro-environment ^6^. In addition, Yuan, et al identified the tumor-promoting role of *PSMB1* in esophageal cancer ^9^. Furthermore, some studies have identified the predictive role of *PSMBs* for cancer drug resistance and immune infiltration. For example, *PSMB8* was found to be related to resistance to Trastuzumab in gastric cancer ^10^. A transcriptional analysis also identified *PSMB2* and *PSMB8* as potential participators in T cell activation ^11^, which suggests the potential application of *PSMBs* as predictors or treatment targets in cancer immunotherapy.

However, whether *PSMBs* could be utilized as an effective biomarker for ccRCC remains elusive. Here we analysed the expression of *PSMBs* in ccRCC and its correlation with prognostic parameters in detail in order to evaluate the potential prognostic value of *PSMBs* for ccRCC.

## 2 Materials and Methods

### 2.1 ONCOMINE

ONCOMINE gene expression array datasets (https://www.oncomine.org, an online cancer microarray database) were used to analyze the transcription levels of *PSMBs* in different cancers. The mRNA expressions of *PSMBs* in clinical cancer specimens were compared with those in normal controls, using a Student’s t test to generate a p value. The cutoffs of p value and fold change were defined as 0.05 and 1.5, respectively.

### 2.2 UALCAN

UALCAN (http://ualcan.path.uab.edu/analysis.html), a comprehensive web resource, provides analyses based on The Cancer Genome Atlas (TCGA) and MET500 cohort data ^12^. In our study, UALCAN was used to analyze the mRNA expressions of *PSMBs* in ccRCC tissues and their association with clinicopathologic parameters. Difference of transcriptional expression was compared by students’ t test and p <0.05 was considered as statically significant.

### 2.3 Human Protein Atlas

The Human Protein Atlas (https://www.proteinatlas.org) is a website that contains immunohistochemistry-based expression data for near 20 highly common kinds of cancers and each tumor type includes 12 individual tumors ^13^. Users can identify tumor-type specific proteins expression patterns that are differentially expressed in a given tumors of type. In this study, direct comparison of protein expression of different *PSMBs* between human normal and ccRCC tissues was performed by immunohistochemistry image.

### 2.4 The Kaplan-Meier Plotter Analysis

The prognostic value of signal transducer and activator of transcription (STAT) mRNA expression was evaluated using an online database, Kaplan-Meier Plotter ^14^, which contained gene expression data and survival information of ccRCC patients. To analyze the overall survival (OS) of patients with ccRCC, patient samples were split into two groups by median expression (high versus low expression) and assessed by a Kaplan-Meier survival plot, with the hazard ratio (HR) with 95% confidence intervals (CIs) and log rank p value.

### 2.5 TCGA and cBioPortal

cBioPortal (www.cbioportal.org), a comprehensive web resource, can visualize and analyze multidimensional cancer genomics data ^15^. Based on TCGA database, genetic alterations, coexpression, and the network module of *PSMBs* was obtained from cBioPortal.

### 2.6 Metascape

Metascape (http://metascape.org) is a reliable, intuitive tool for gene annotation, and gene list enrichment analysis ^16^. Based on the functional annotation of gene/protein lists, Metascape can facilitate data-driven decisions. In this study, the “Express Analysis” module was used to further verify the enrichment of *PSMBs* and closely related neighbor genes.

### 2.7 GeneMANIA

GeneMANIA (http://genemania.org) is a flexible user-friendly web site for generating hypotheses about gene function, analyzing gene lists and prioritizing genes for functional assays ^17^. In this study, GeneMANIA was use to analyse differential expressed *PSMBs*.

### 2.8 STRING

STRING (https://string-db.org/) aims to collect, score, and integrate all publicly available sources of protein–protein interaction (PPI) data, and to complement these with computational predictions of potential functions ^18^. We conducted a PPI network analysis of differentially expressed *PSMBs* to explore the interactions among them with STRING.

### 2.9 DAVID6.8

DAVID6.8 (https://david.ncifcrf.gov/home.jsp) is a comprehensive, functional annotation website that helps investigators better clarify the biological function of submitted genes ^19^. In our study, the Gene Ontology (GO) enrichment analysis and Kyoto Encyclopedia of Genes and Genomes (KEGG) pathway enrichment analysis of *PSMBs* and closely related neighbor genes were isolated from DAVID6.8 and visualized with R project using a “ggplot2” package and a p < 0.05. Biological processes (BP), cellular components (CC), and molecular function (MF) were included in the GO enrichment analysis.

### 2.10 TIMER

TIMER (https://cistrome.shinyapps.io/timer/) is a reliable, intuitive tool that provides systematic evaluations of the infiltration of different immune cells and their clinical impact ^20^. In our study, “Gene module” was used to evaluate the correlation between the level of *PSMBs* expression and the infiltration of immune cells. “Survival module” was used to evaluate the correlation among clinical outcome and the infiltration of immune cells and *PSMBs* expression.

## 3 Results

### 3.1 Transcription Levels of *PSMBs* in Patients with ccRCC

To date, 10 different members of the *PSMBs* have been identified in mammalian cells. It was the first time that mRNA expressions of *PSMBs* were analyzed and compared to normal tissues by Oncomine database (**Figure** 1 and **Table 1**). Oncomine analysis revealed that the mRNA expressions of *PSMB2*, *PSMB7*, *PSMB8*, *PSMB9*, and *PSMB10* were upregulated in ccRCC versus normal renal tissue (**Figure 1**).

**Figure 1.**
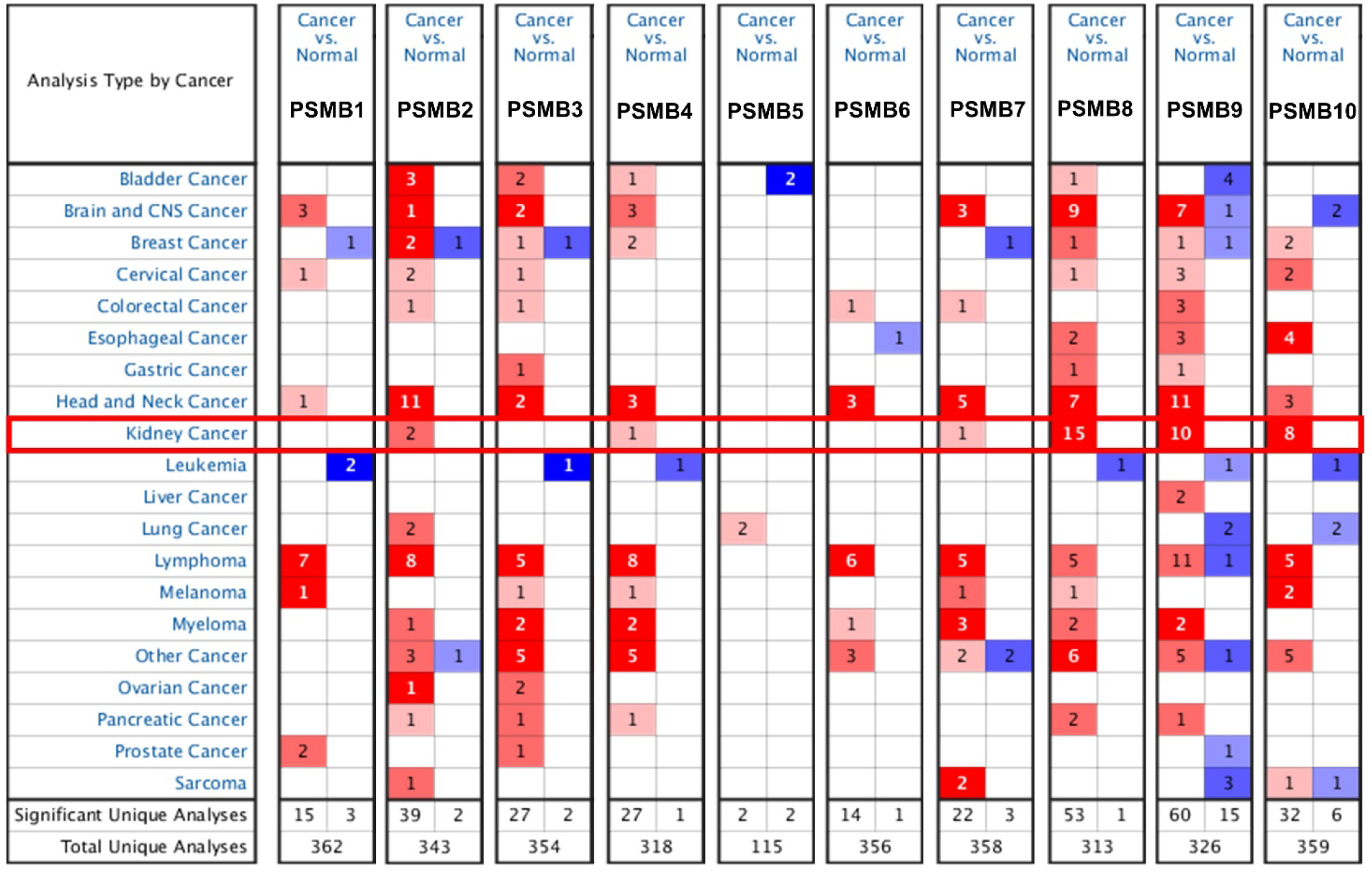
mRNA levels of *PSMBs* in ccRCC (ONCOMINE). The figure shows the numbers of datasets with statistically significant mRNA overexpression (red) or downregulated expression (blue) of *PSMBs*.

**Table 1.** The mRNA levels of PSMBs in different types of RCC tissues and normal renal tissues at transcriptome level (ONCOMINE).

|  | Types of kidney cancer vs. Normal | Fold change | P-value | t-test | References |
| --- | --- | --- | --- | --- | --- |
| *PSMB2* | Renal Wilms Tumor vs. Normal | 2.099 | 0.006 | 4.104 | Yusenko Renal |
| *PSMB7* | Renal Wilms Tumor vs. Normal | 2.312 | 0.004 | 3.930 | Yusenko Renal |
| *PSMB8* | Clear Cell Renal Cell Carcinoma vs. Normal | 2.980 | 2.37E-8 | 11.319 | Lenburg Renal |
|  | Clear Cell Renal Cell Carcinoma vs. Normal | 2.765 | 2.92E-9 | 15.193 | Higgins Renal |
|  | Clear Cell Renal Cell Carcinoma vs. Normal | 15.139 | 1.08E-6 | 8.416 | Gumz Renal |
|  | Clear Cell Renal Cell Carcinoma vs. Normal | 9.429 | 6.91E-5 | 11.121 | Yusenko Renal |
|  | Clear Cell Renal Cell Carcinoma vs. Normal | 2.995 | 1.08E-8 | 7.805 | Jones Renal |
| *PSMB9* | Clear Cell Renal Cell Carcinoma vs. Normal | 5.044 | 3.11E-9 | 11.645 | Lenburg Renal |
|  | Clear Cell Renal Cell Carcinoma vs. Normal | 4.688 | 1.88E-8 | 10.627 | Gumz Renal |
|  | Clear Cell Renal Cell Carcinoma vs. Normal | 5.267 | 1.70E-14 | 12.295 | Jones Renal |
|  | Clear Cell Renal Cell Carcinoma vs. Normal | 12.722 | 7.25E-4 | 6.878 | Yusenko Renal |
| *PSMB10* | Clear Cell Renal Cell Carcinoma vs. Normal | 6.000 | 5.74E-7 | 11.706 | Yusenko Renal |
|  | Clear Cell Renal Cell Carcinoma vs. Normal | 2.330 | 4.24E-4 | 4.098 | Lenburg Renal |
|  | Clear Cell Renal Cell Carcinoma vs. Normal | 2.595 | 5.49E-10 | 9.048 | Jones Renal |
|  | Clear Cell Renal Cell Carcinoma vs. Normal | 2.829 | 6.21E-5 | 4.886 | Gumz Renal |

**Table 2.** The GO function enrichment analysis of PSMB family members in ccRCC.

| GO | Category | Description |
| --- | --- | --- |
| GO:0002479 | GO biological processes | antigen processing and presentation of exogenous peptide antigen via MHC class I |
| GO:0051437 | GO biological processes | positive regulation of ubiquitin-protein ligase activity involved in regulation of mitotic cell cycle transition |
| GO:0031145 | GO biological processes | anaphase-promoting complex-dependent catabolic process |
| GO:0051436 | GO biological processes | negative regulation of ubiquitin-protein ligase activity involved in mitotic cell cycle |
| GO:0000502 | GO cellular components | proteasome complex |
| GO:0005839 | GO cellular components | proteasome core complex |
| GO:0005654 | GO cellular components | nucleoplasm |
| GO:0005829 | GO cellular components | cytosol |
| GO:0005634 | GO cellular components | nucleus |
| GO:0004298 | GO molecular functions | threonine-type endopeptidase activity |
| GO:0044822 | GO molecular functions | poly(A) RNA binding |
| GO:0005515 | GO molecular functions | protein binding |
| GO:0003735 | GO molecular functions | structural constituent of ribosome |

*PSMB2* was over expressed in renal wilms tumor (fold change = 2.099) compared to normal sample in Yusenko’s dataset (**Table 1**). The over-expression of *PSMB7* was also explored in renal wilms tumor with a fold change of 2.312. *PSMB8*was found to be higher expressed in ccRCC versus normal samples in five datasets, which are Lenburg’s dataset (fold change = 2.980), Higgins’s dataset (fold change = 2.765), Gumz’s dataset (fold change = 15.139), Yusenko’s dataset (fold change = 9.429) and Jones’s dataset (fold change = 2.995) (**Table 1**). In the contrast between ccRCC and normal tissues (**Table 1**), we found *PSMB9* over-expression in Lenburg’s dataset (fold change = 5.044), Gumz’s dataset (fold change = 4.688), Jones’s dataset (fold change = 5.267) and Yusenko’s dataset (fold change = 12.722). *PSMB10* was also found to be over expressed in ccRCC compared with normal samples in Yusenko’s dataset (fold change = 6.000), Lenburg’s dataset (fold change = 2.330), Jones’s dataset (fold change = 2.595) and Gumz’s dataset (fold change = 2.829) (**Table 1**).

### 3.2 Relationship Between the mRNA Levels of *PSMBs* and the Clinico-pathological Parameters of Patients with ccRCC

We compared the mRNA expression of *PSMBs* between ccRCC and normal tissues with UALCAN (http://ualcan.path.uab.edu) (**Figure 2**). The results showed that the expression levels of *PSMB1/2/4/7/8/9/10* were higher in ccRCC tissues than in normal tissues, whereas the expression levels of *PSMB5/6* were significantly reduced. Besides, the expression level of *PSMB3* was similar in tumor tissues and normal samples.

**Figure 2.**
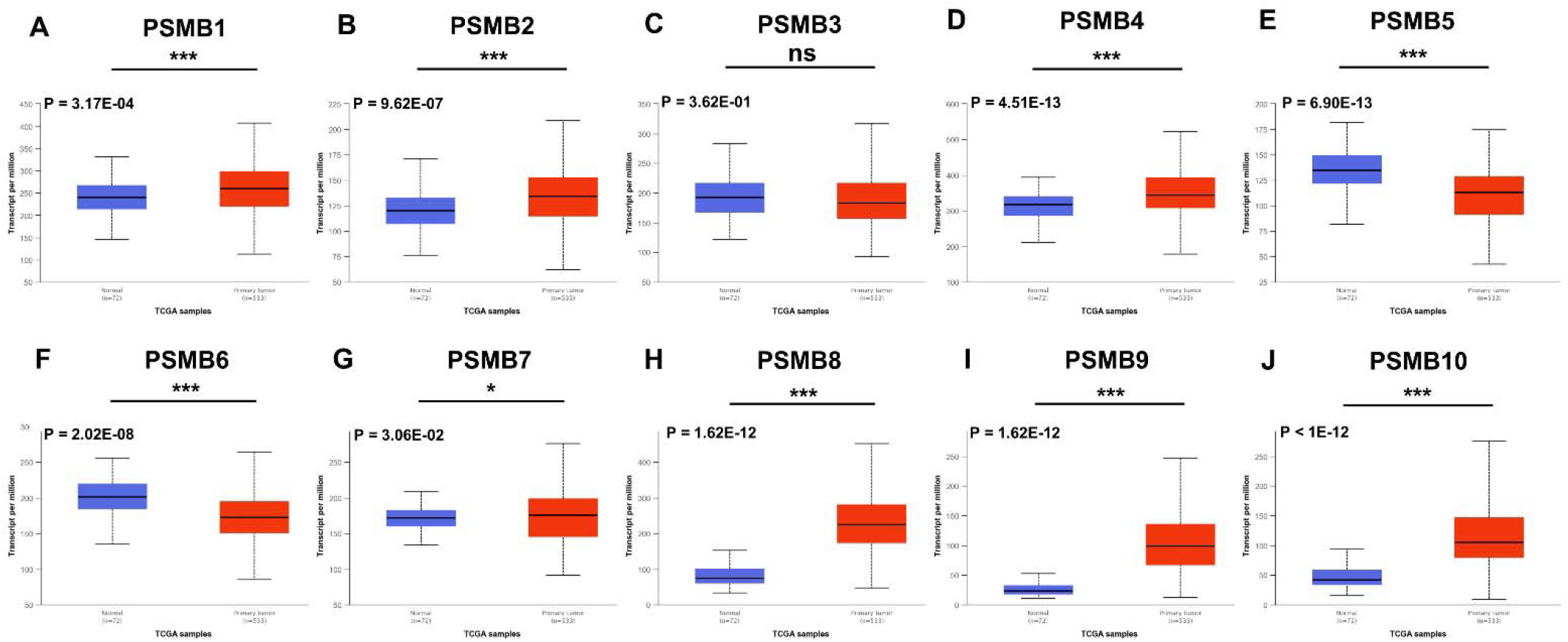
The transcription of *PSMBs* in ccRCC (UALCAN). The transcriptional levels of (A) *PSMB1*, (B) *PSMB2*, (D) *PSMB4*, (G) *PSMB7*, (H) *PSMB8*, (I) *PSMB9* and (J) *PSMB10* in RCC tissues were significantly elevated while the transcriptional levels of (E) *PSMB5*, (F) *PSMB6*, were significantly reduced. The P value was set at 0.05. * p < 0.05, ** p < 0.01, *** p < 0.001.

Then we explored the relationship between mRNA expression levels of *PSMBs* with clinico-pathological parameters of ccRCC patients by UALCAN, including patients’ individual cancer stages and tumor grades. We found higher expression levels of *PSMB1/2/3/4/7/8/9/10* in more advanced cancer stages (**Figure 3**). However lower expressions of *PSMB5* and *PSMB6* were found in more advanced stages (**Figure 3**). The highest mRNA expressions of *PSMB1/2/3/4/7/8/9/10* were found in stage 4 (**Figure 3A-D, G-J**), while the highest mRNA expressions of *PSMB5* and *PSMB6* were found in stage 1 (**Figure 3E, F**).

**Figure 3.**
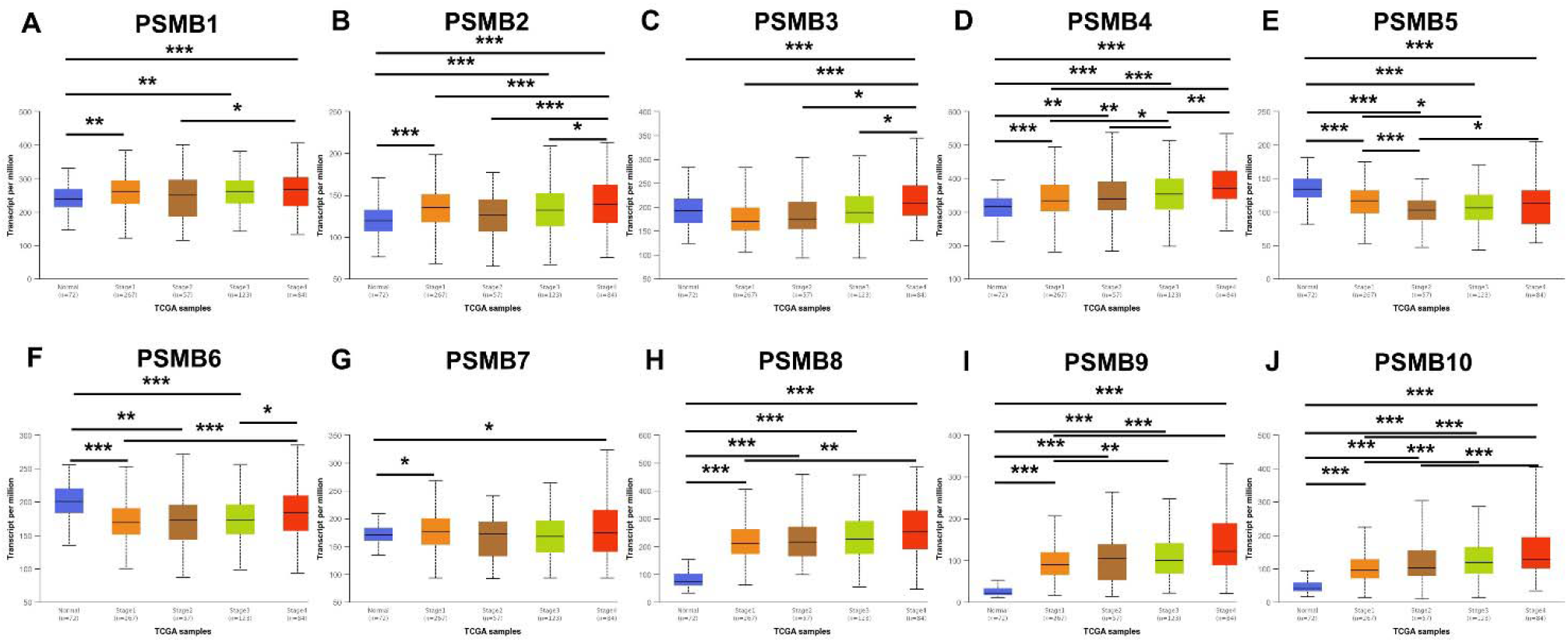
Correlation between mRNA expression of *PSMBs* and pathological stage of ccRCC patients (UALCAN). (A) *PSMB1*, (B) *PSMB2*, (C) *PSMB3*, (D) *PSMB4*, (E) *PSMB5, (F)PSMB6, (G)PSMB7*, (H) *PSMB8*, (I) *PSMB9*, (J) *PSMB10*. The p value was set at 0.05. * p < 0.05, ** p < 0.01, *** p < 0.001.

Similarly, as was shown in **Figure 4**, mRNA expression levels of *PSMBs* were associated with tumor grades. As tumor grade increased, the mRNA expression of *PSMB1/2/3/4/7/8/9/10* tended to be higher. The highest expression levels of *PSMB1/2/3/4/8/9/10* were found in tumor grade 4 (**Figure 4A-D, H-J**), while the highest mRNA expression of *PSMB7* was found in grade 2 (**Figure 4G**). However, the highest mRNA expression levels of *PSMB5/6* were found in grade 1, and as tumor grade increased, the mRNA expression of *PSMB5/6* tended to be lower (**Figure 4E, F**).

**Figure 4.**
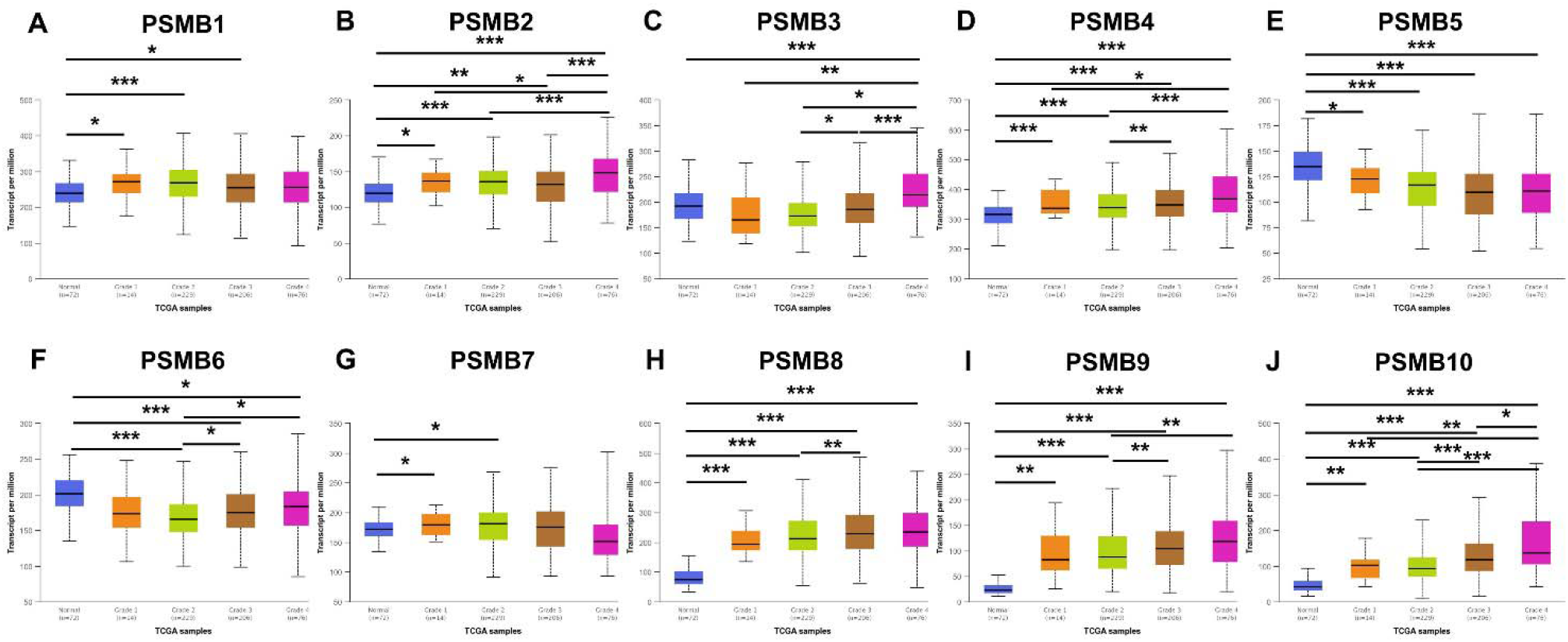
The association of mRNA expression of *PSMBs* with tumor grades of ccRCC patients (UALCAN). (A) *PSMB1*, (B) *PSMB2*, (C) *PSMB3*, (D) *PSMB4*, (E) *PSMB5, (F)PSMB6, (G)PSMB7*, (H) *PSMB8*, (I) *PSMB9*, (J) *PSMB10*. The p value was set at 0.05. * p < 0.05, ** p < 0.01, *** p < 0.001.

Next, we explored the protein expression of *PSMBs* in ccRCC patients by the Human Protein Atlas. *PSMB3/4/8/9/10* proteins are not detected or medium expressed in normal tissues, while high expressions of them were found in ccRCC tissues (**Figure 5C-D, H-J**). Besides, medium protein expression of *PSMB5* was found in normal samples, whereas it was not detected in cancer tissues (**Figure 5E**). However, high protein expressions of *PSMB1/2/7* (**Figure 5A, B, G**) and medium expression of *PSMB6* (**Figure 5F**) were observed both at normal tissues and ccRCC tissues.

**Figure 5.**
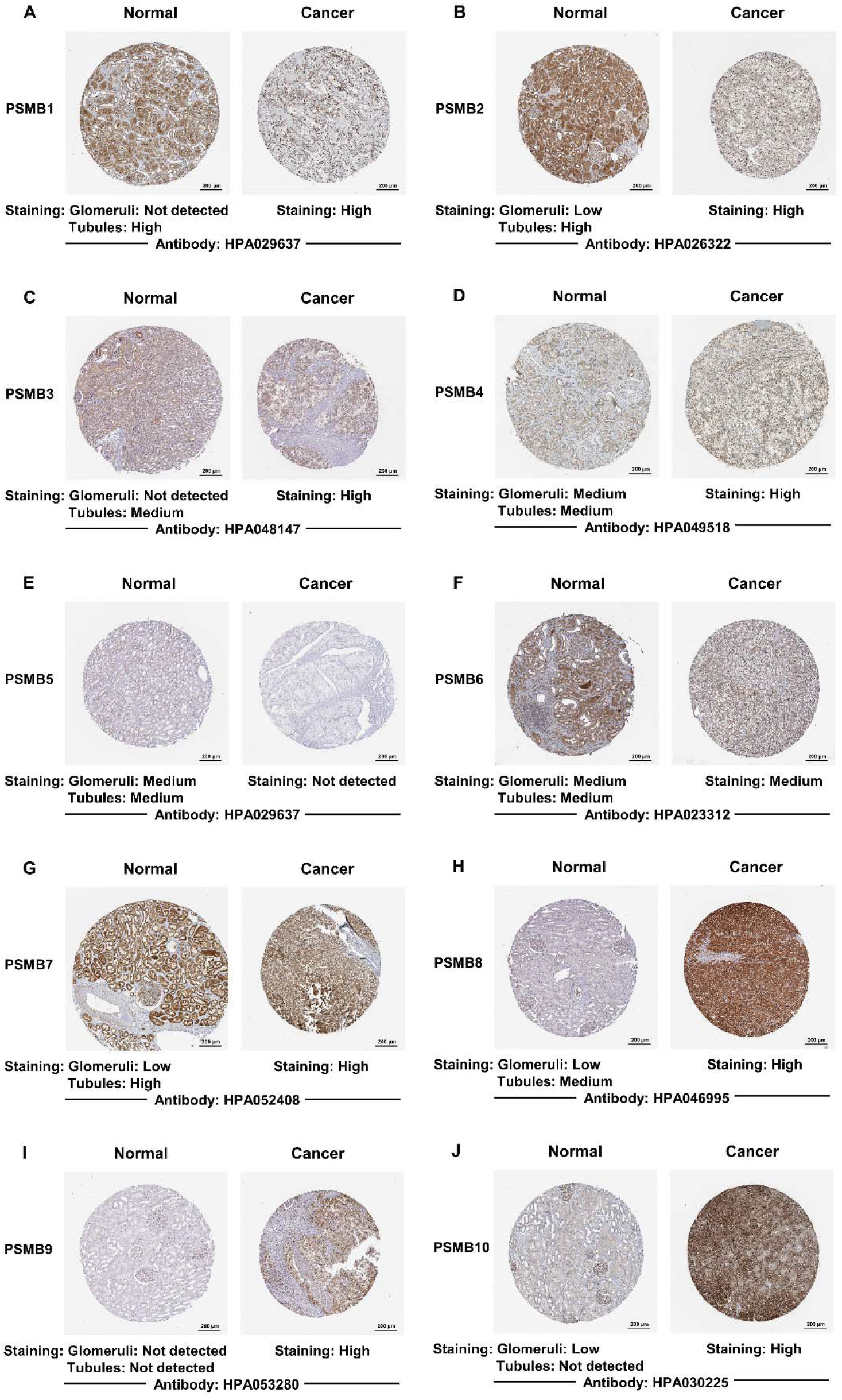
Representative immunohistochemistry images of *PSMBs* in ccRCC tissues and normal renal tissues (Human Protein Atlas). (A) *PSMB1*, (B) *PSMB2*, (C) *PSMB3*, (D) *PSMB4*, (E) *PSMB5*, (F)*PSMB6*, (G)*PSMB7*, (H) *PSMB8*, (I) *PSMB9*, (J) *PSMB10*.

### 3.3 Prognostic Values of *PSMBs* in Patients With ccRCC

We explored the prognostic significance of *PSMBs* in ccRCC patients by using Kaplan-Meier plotter. We found that higher mRNA expression level of *PSMB1* (p = 3e-04), *PSMB2* (p = 7.1e-05), *PSMB3* (p = 1.3e-06), *PSMB4* (p = 1.1e-05), *PSMB5* (p = 0.0091), *PSMB6* (p = 0.00074) and *PSMB10* (p = 0.0036) were significantly associated with shorter OS of patients with ccRCC (**Figure 6A-E, F and J**), whereas, higher mRNA expression level of *PSMB7* (**Figure 6G**) was connected to favorable OS of patients with ccRCC (p = 0.00035).

**Figure 6.**
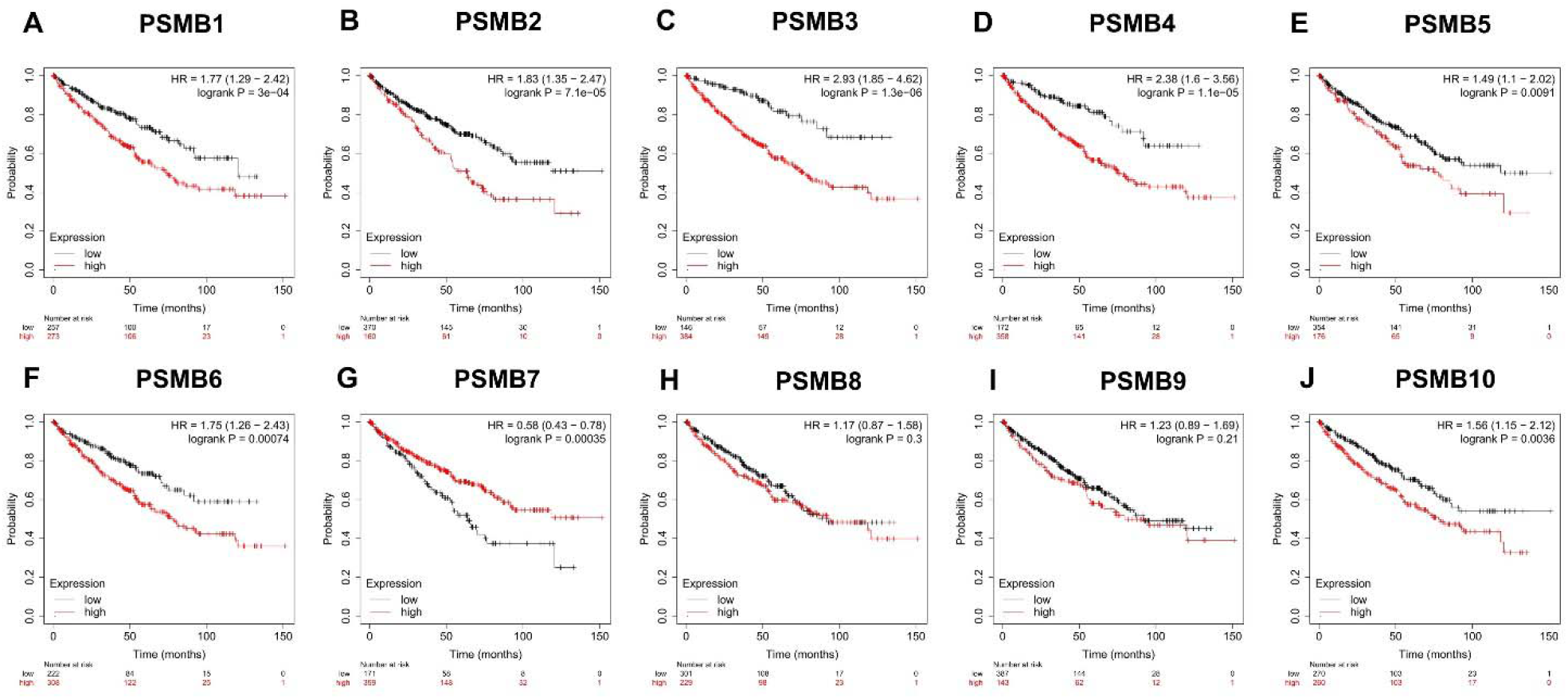
The prognostic value of *PSMBs* in ccRCC patients in overall survival curve (Kaplan-Merier Plotter). (A) *PSMB1*, (B) *PSMB2*, (C) *PSMB3*, (D) *PSMB4*, (E) *PSMB5*, (F)*PSMB6*, (G)*PSMB7*, (H) *PSMB8*, (I) *PSMB9*, (J) *PSMB10*.

### 3.4 Genetic Alteration, Co-expression, Neighbor Gene Network, and Interaction Analyses of *PSMBs* in Patients With ccRCC

We used cBioPortal to analyze the genetic alteration of *PSMBs* and the correlation of *PSMBs* with each other (Provisional datasets of TCGA). We found that the up-regulated PSMBs mRNA accounted for the majority of alteration frequency relatively, followed by the down-regulated *PSMBs* mRNA (**Figure 7A**). *PSMB1* had the highest percentage of genetic alterations (10%, **Figure 7A**). The next few with higher genetic changes were *PSMB6* (9%, **Figure 7A**), *PSMB4* (8%, **Figure 7A**) and *PSMB5* (7%, **Figure 7A**). Results of Pearson correlation analysis suggested positive correlations between *PSMBs*, including *PSMB1* and *PSMB6, PSMB1* and *PSMB7, PSMB3* and *PSMB6, PSMB3* and *PSMB10, PSMB4* and *PSMB6, PSMB6* and *PSMB7*. We also found significant positive correlation between *PSMB8* and *PSMB9* and *PSMB10*. Besides, negative associations between *PSMB5* and *PSMB8, PSMB5* and *PSMB9, PSMB5* and *PSMB10* were explored.

**Figure 7.**
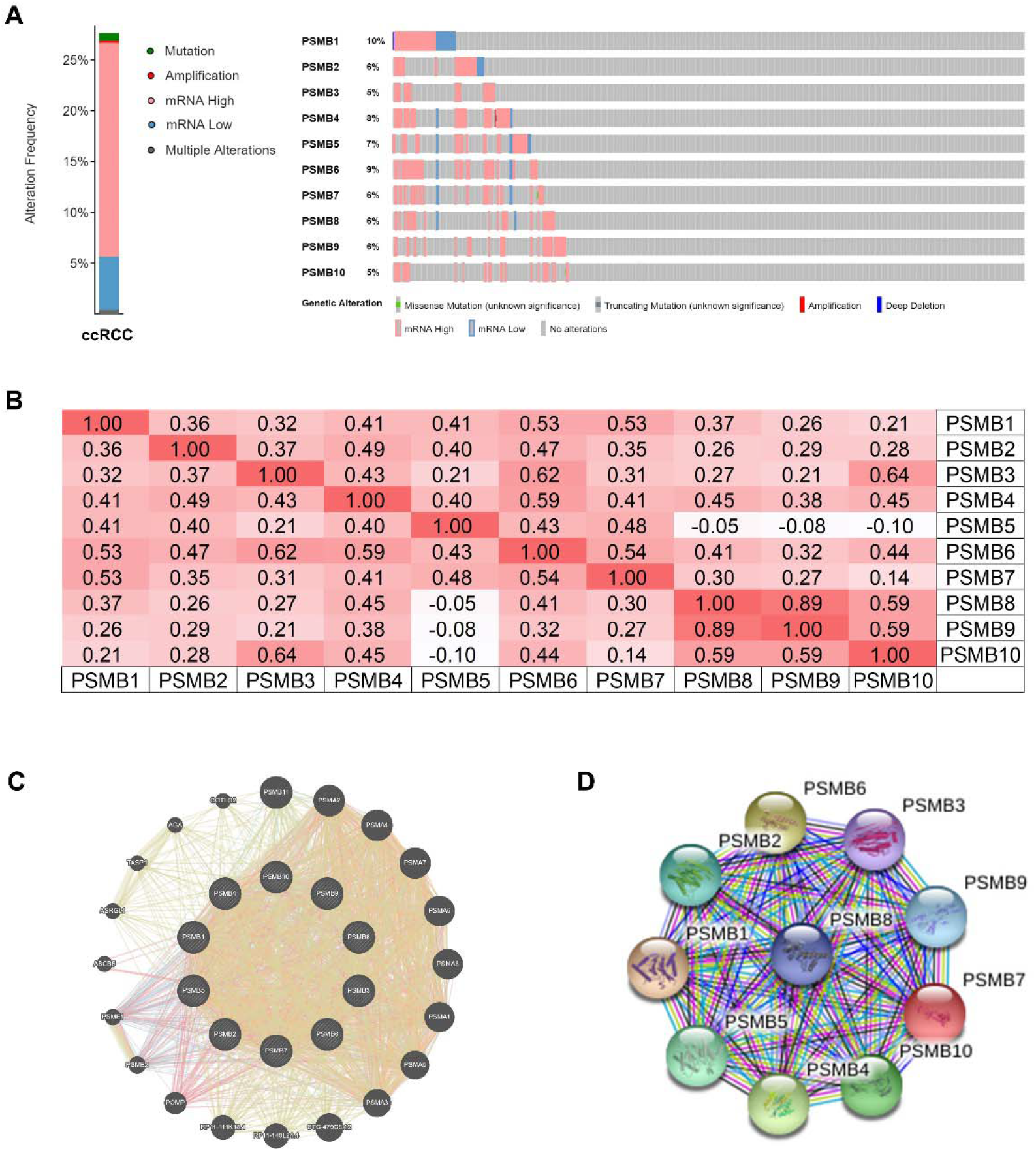
Genetic alteration, neighbor gene network, and interaction analyses of different expressed *PSMB* family members in ccRCC patients. (A) *PSMBs* gene expression and mutation analysis in ccRCC (cBioPortal). (B) Correction between different *PSMBs* in ccRCC (cBioPortal). (C, D) Protein–protein interaction network of different expressed *PSMBs* (GeneMANIA, STRING).

Results of GeneMANIA also revealed that the functions of differential expressed *PSMBs (PSMB1*, *PSMB2*, *PSMB3*, *PSMB4*, *PSMB5*, *PSMB6*, *PSMB7*, *PSMB8*, *PSMB9*, and *PSMB10*) were primarily related to the regulation of cellular processes (**Figure 7C**). Moreover, we conducted a protein protein interaction (PPI) network analysis of differentially expressed *PSMBs* with STRING to explore the potential interactions among them. As expected, several nodes of 12 and several edges of 45 were obtained in the PPI network (**Figure 7D**). The function of these differentially expressed *PSMB* members was associated with the deubiquitinating of protein and post-translational protein modification.

### 3.5 Functional Enrichment Analysis of PSMBs in Patients With ccRCC

We used GO and KEGG to explore the possible functions of the *PSMBs* and their adjacent genes. The obvious biological processes (**Figure 8A**) involved in these genes are GO:0002479 (antigen processing and presentation of exogenous peptide antigen via MHC class I), GO:0051437 (positive regulation of ubiquitin-protein ligase activity involved in regulation of mitotic cell cycle transition), GO:0031145 (anaphase-promoting complex-dependent catabolic process) and GO:0051436 (negative regulation of ubiquitin-protein ligase activity involved in mitotic cell cycle). We also found several highly enriched items in cellular component group (**Figure 8B**) such as GO:0000502 (proteasome complex), GO:0005839 (proteasome core complex), GO:0005654 (nucleoplasm), GO:0005829 (cytosol) and GO:0005634 (nucleus). In addition, we found that the molecular functions (**Figure 8C**) associated with these genes significantly were GO:0004298 (threonine-type endopeptidase activity), GO:0044822 (poly(A) RNA binding), GO:0005515 (protein binding) and GO:0003735 (structural constituent of ribosome). Subsequently, KEGG analysis (**Figure 8D**) was performed to identify pathways that were significantly associated with these genes, including hsa03050 (Proteasome), hsa04612 (Antigen processing and presentation), hsa03040 (Spliceosome), hsa05169 (Epstein-Barr virus infection), hsa03013 (RNA transport), hsa05168 (Herpes simplex infection) and hsa03010 (Ribosome).

**Figure 8.**
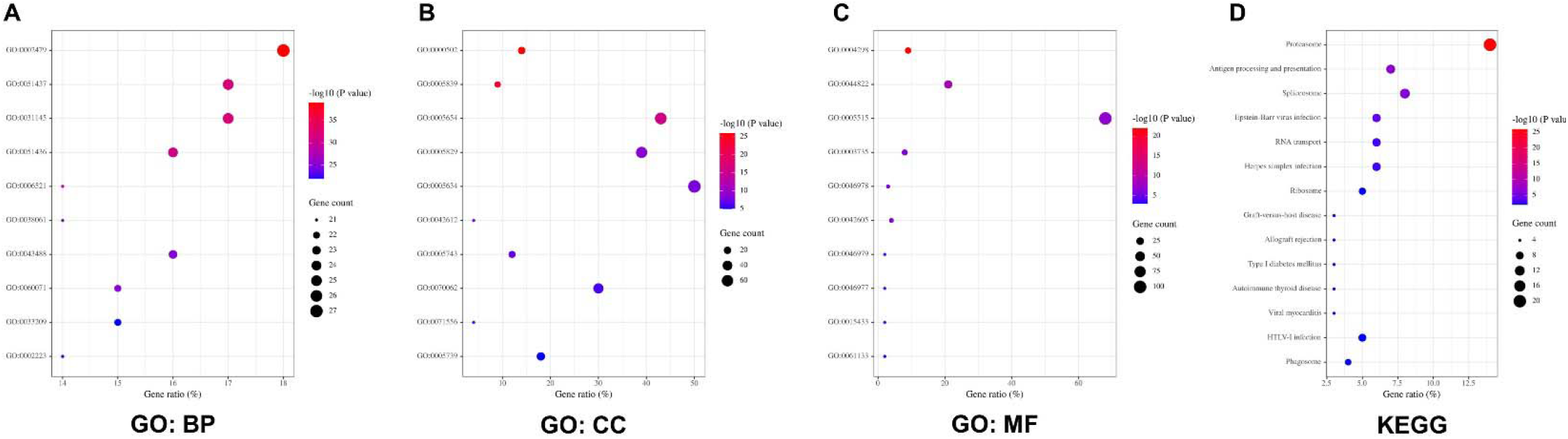
The functions of *PSMBs* and genes significantly associated with *PSMBs* alterations in ccRCC. The functions of *PSMBs* and genes significantly associated with *PSMBs* alterations were predicted by the analysis of gene ontology (GO) by (Database for Annotation, Visualization and Integrated Discovery) tools. GO enrichment analysis predicted the functional roles of target host genes based on three aspects, including (A) biological processes, (B) cellular components, and (C) molecular functions. The functions of *PSMBs* and genes significantly associated with *PSMBs* alterations were predicted by the analysis of Kyoto Encyclopedia of Genes and Genomes (KEGG) by DAVID tools (D).

### 3.6 Immune Cell Infiltration of PSMBs in Patients With ccRCC

As reported, *PSMB8* and *PSMB9* were related to immune system. Therefore, we deeply explored the correlation between the mRNA expression levels of *PSMBs* and immune cell infiltration by TIMER database. We found a positive correlation between expression level of *PSMB1* and infiltration level of CD8+ T cells (Cor = 0.237, p = 5.32e-7), and a negative correlation between expression level of *PSMB1* and infiltration level of CD4+ T cells (Cor = −0.112, p = 1.65e-2; **Figure 9A**). Expression level of *PSMB2* was significantly positive connected with infiltration level of B cells, CD8+ T cells, CD4+ T cells, macrophages, neutrophils and dendritic cells with all p < 0.01 (**Figure 9B**). There was a negative correlation between expression of *PSMB3* and the infiltration of CD4+ T cells (Cor = −0.207, p = 7.51e-6), macrophages (Cor = −0.268, p = 8.00e-9) and neutrophils (Cor = −0.209, p = 6.63e-6; **Figure 9C**). Expression of *PSMB4* was also negatively related to infiltration level of CD4+ T cells (Cor = −0.203, p = 1.13e-5) and macrophages (Cor = −0.165, p = 4.67e-4; **Figure 9D**). The expression level of *PSMB5* was not correlated with the infiltration of these immune cells (**Figure 9E**). Expression level of *PSMB6* was found positively associated with infiltration level of B cells (Cor = 0.096, p = 4.08e-2) and negatively associated with infiltration level of CD4+ T cells (Cor = −0.146, p = 1.75e-3), macrophages (Cor = −0.231, p = 8.13e-7) and neutrophils (Cor = −0.193, p = 3.20e-5; **Figure 9F**). Similarly, expression of *PSMB7* has positive connected with the infiltration of CD8+ T cells (Cor = 0.141, p = 3.09e-3) and negative connected with the infiltration of CD4+ T cells (Cor = −0.164, p = 4.10e-4), macrophages (Cor = −0.131, p = 5.60e-3) and neutrophils (Cor = −0.17, p = 2.49e-4; **Figure 9G**). Moreover, we found that *PSMB8, PSMB9* and *PSMB10* shared similar characteristic in the correlation between expression level of them and infiltration level of immune cells. To be specific, significantly positive correlation between the expression of *PSMB8, PSMB9* and *PSMB10* and the infiltration level of B cells, CD8+ T cells, neutrophils and dendritic cells was explored, with all p < 0.01 (**Figure 9H-J**).

**Figure 9.**
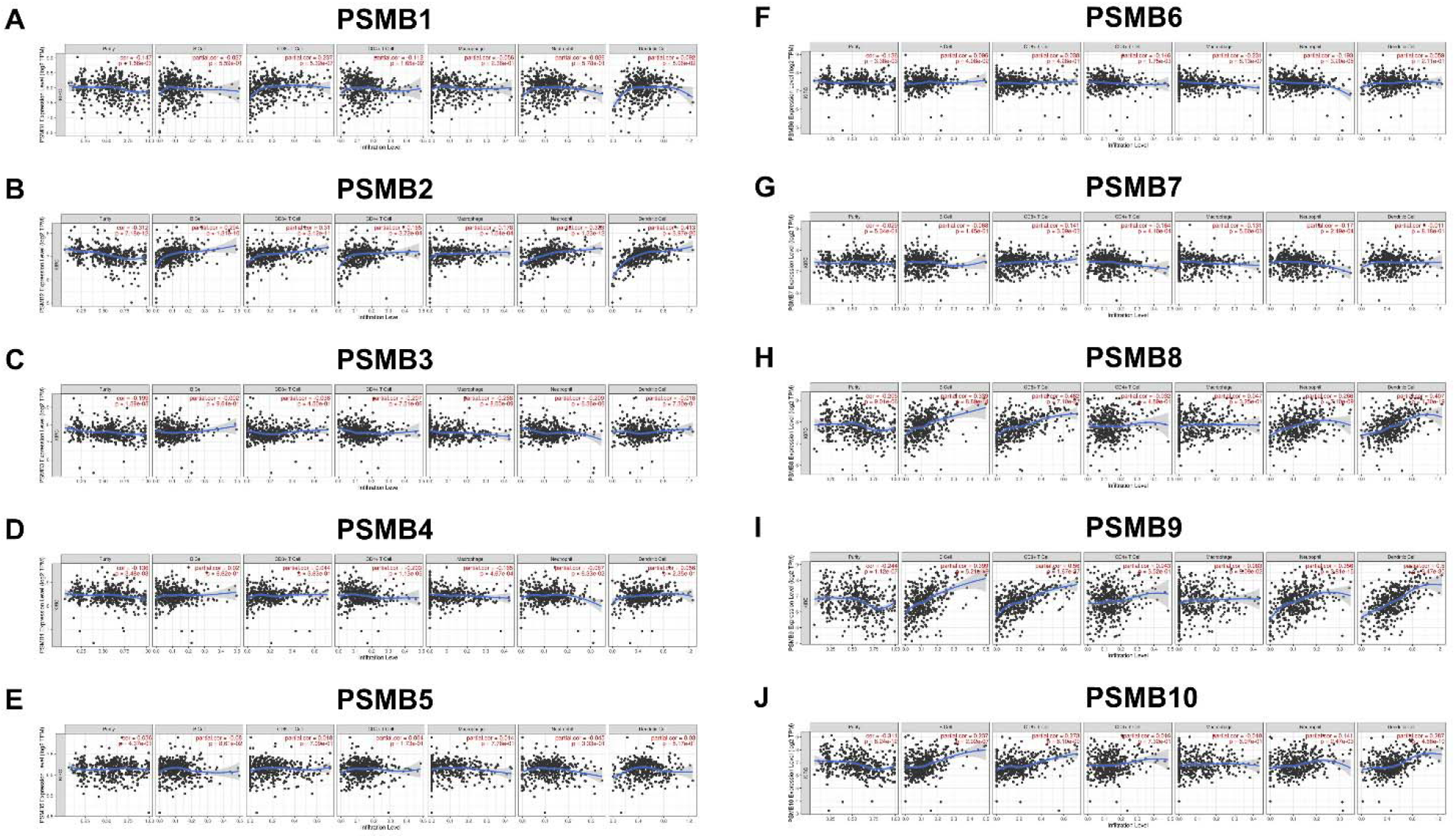
Correlation between different expressed *PSMBs* and immune cells infiltration (TIMER). (A) *PSMB1*, (B) *PSMB2*, (C) *PSMB3*, (D) *PSMB4*, (E) *PSMB5, (F)PSMB6, (G)PSMB7*, (H) *PSMB8*,(I) *PSMB9*, (J) *PSMB10*.

## 4 Discussion

To summarize, we found the expressions of a series of *PSMB* members, including *PSMB2/7/8/9/10*,were significantly increased in ccRCC, and the expression of *PSMB1/2/3/4/8/9/10* were related to advanced tumor stage and grade. Moreover, the higher mRNA expression of *PSMB1/2/3/4/5/6/10* were associated with a shorter OS. Furthermore, the protein expression level of *PSMB1/2/3/4/7/8/9/10* in ccRCC was higher than normal renal tissues. This study indicated potential roles of *PSMBs* as a useful biomarker in the diagnosis, prognosis prediction and treatment guidance of ccRCC.

Proteasome, which is mainly encoded by *PSMBs*, mainly participated in the process of degrading poly-ubiquitinated intracellular proteins to remove unfolded or misfolded proteins. In addition, proteasome also regulate other multiple cellular processes. Enhanced proteasome activity was reported to be detected in different cancers ^21^. *PSMB1* is a member of proteasome β family and its exact role played in cancer has not been elucidated before. In our study, *PSMB1* is highly expressed in ccRCC tissues compared with normal tissues. Higher transcriptional expression of *PSMB1* was correlated with a higher malignant degree and worse prognosis of ccRCC patients, which indicated an oncogene role of *PSMB1* in ccRCC. However, the exact mechanism is still elusive. Gergely Varga, et al found a suboptimal response to bortezomib treatment in myeloma could be caused by a *PSMB1* minor allele mutation, which suggested that the loss of proteasome activity of *PSMB1* may decrease sensitivity to bortezomib treatment by a large supply of undegraded proteins ^22^. Another study also found good response to bortezomib is relying on high expression of *PSMB1*^23^. Recently, with the rapid progress of immune-checkpoint inhibitor therapy for cancer, much more attention is being attracted to the immune status in the tumor micro-environment, including ccRCC. Wu, et al demonstrated that *PSMB1* initiated a negative regulatory role in the innate antiviral immunity by inhibiting TLR3 and RLR pathway ^24^. These findings reminded us that proteasome activity may be a sword with two sides in tumorigenesis. Although its higher activity may predict worse survival, loss of *PSMBs* function may also provide potential escape pathway for tumor cell when treated with targeted therapy. Except for the classic role of *PSMBs* in protein degradation, it also participates in the process of cancer progression through cell-cycle pathway. Yuan, et al proposed that proteasomal β1 subunit promoted tumorigenesis in different malignancies by degrading p27^Kip1^, which is a crucial regulator of cell-cycle ^9^, which may also give explanations for the oncogene function of *PSMBs* in ccRCC.

Although *PSMB2* has been found to play major roles in multiple cancers, including hepatocellular carcinoma^25^, Pancreatic ductal adenocarcinoma^26^ and breast cancer^27^. Until now, little was known about the expression and role of *PSMB2* in ccRCC. In our study, we found that the expression level of *PSMB2* in ccRCC tissues was higher than that in normal tissues. In addition, high expression level of *PSMB2* was correlated to advanced tumor stage and increased tumor grades in patients with ccRCC. Similar results could be detected from *PSMB3*, *PSMB4*, *PSMB7 and PSMB10*. High *PSMB2* expression was significantly connected to shorter OS and higher infiltration level of B cells, CD8+ T cells, CD4+ T cells, macrophages, neutrophils and dendritic cells in all of the patients with ccRCC.

Accoding to the analysis of *PSMB3* by CancerSEA (http://biocc.hrbmu.edu.cn/CancerSEA), *PSMB3* was known to play a role in controlling cell cycle progression. As reported, down-regulation of *PSMB3* were revealed as trisomy-associated alterations involved in regulating genome stability, which related to the global proteome equilibrium of colorectal cancer cells^28^. In this study, the expression of *PSMB3* in ccRCC tissues was higher than that in normal tissues. We also found that *PSMB3* expression was significantly correlated with advanced tumor stage in patients with ccRCC and highest expression levels of *PSMB3* was found in tumor grade4. Further, analysis revealed that *PSMB3* overexpression was associated with reduced OS in ccRCC patients. In addition, there was a negative correlation between expression of *PSMB3* and the infiltration of CD4+ T cells, macrophages and neutrophils.

*PSMB4* is an important member of *PSMB* family, which is elevated in a variety of malignancies ^29^. In this study, the expression level of *PSMB4* in ccRCC tissues was higher than that in normal tissues. Besides, high expression level of *PSMB4* was correlated to advanced tumor stage in patients with ccRCC. Higher *PSMB4* expression was significantly connected to shorter OS and higher infiltration level of B cells, CD8+ T cells, CD4+ T cells, macrophages, neutrophils and dendritic cells in all of the patients with ccRCC.

Intriguingly, different from other *PSMBs*, a relatively lower mRNA and protein expression of *PSMB5* and *PSMB6* was found in ccRCC tumor tissues compared with normal tissues. Wang, et al recognized *PSMB5* as a strong immune suppressor in breast cancer, functioning by promoting monocytes to differentiate into M2 macrophages ^6^. More recently, Guo, et al found that *PSMB5* was modulated by Tribbles homolog 2 (TRIB2), which helped reduce the ubiquitin proteasome system activity and protect the liver cancer cells from oxidative damage ^30^. These results suggested that PSMB5 may also play important roles in the development of ccRCC. However, the low expression on ccRCC tumor tissue my limit its clinical practice.

With regards to *PSMB6*, current investigations are lacked. Although lower expression of *PSMB6* was found in cancer tissues on both mRNA and protein levels, prognostic analysis indicated that *PSMB6* may also exert pro-tumor roles in the progression of ccRCC, but the definite mechanisms required to be resolved in the future.

Surprisingly, *PSMB7*, which also belongs to proteasome β1 family, predicted a better prognosis of ccRCC patients. On the contrary, G Munkácsy, et al demonstrated that high expression of *PSMB7* represents a bad prognosis under the treatment of doxorubicin and paclitaxel. Nevertheless, the functions of *PSMB7* in the ccRCC development varied dependent on the treatment contexts, which may explain the different relationships between *PSMB7* and malignancies ^31^.

A bioinformatic analysis revealed a range of genes could help identify immune infiltration characterics, including *PSMB8* and *PSMB9* ^32^. By using Timer database, we performed an analysis of the relationship between the expression degree of *PSMBs* and immune infiltration of various immune cells. Higher expression of *PSMB1/3/4/6* was found to be negatively associated with a less infiltration of CD4+ T cells, which indicated that *PSMBs* could participated in the adaptive immune regulation in ccRCC, mainly by inhibiting CD4+ T cell differentiation. However, due to the high heterogeneity of CD4+ T cell, deeper analysis is required. Moreover, Nishida, et al reported a significant up-regulation of CD4+/ CD8+ T cell in the RCC patients and the CD4+/ CD8+ T cells expressed a large amount of immune-checkpoint molecules, including PD-1, TIM-3, OX40 and so on ^33^. In our results, higher expression of *PSMB8/9/10* is strongly correlated with more infiltration of CD8+ T cell. We hypothesize that these CD8+ T cells may represent a large group of exhausted population of T cell in the tumor-microenvironment.

*PSMB10* was also found to be over expressed in ccRCC compared with normal samples. Higher expression level of *PSMB10* was related to advanced stages and increased tumor grades in the patients with ccRCC. As reported, *PSMB10* was a missing gene for proteasome-associated autoinflammatory syndrome, which indicated that *PSMB10* may play a role in immune mechanism ^34^. In our study, we found significantly positive correlation between the expression of *PSMB10* and the infiltration level of B cells, CD8+ T cells, neutrophils and dendritic cells.

In this study, we systemically performed an analysis of the expressive and prognostic value of *PSMBs* for ccRCC for the first time. However, the exact biological roles and concrete mechanisms utilized by *PSMBs* to participate in the pathological process of ccRCC are still elusive, more and deeper studies are required to elucidate the relationship between *PSMBs* and ccRCC and put *PSMBs* in clinical practice as a useful predictive biomarker and therapeutic target.

## 5 Conclusions

We systematically analyzed the expression and prognostic value of PSMBs in ccRCC, and we provided a thorough understanding of the complexity of the molecular biological properties of ccRCC. Our results indicated that the increased expression of *PSMB4* and *PSMB10* in ccRCC tissues might play an important role in ccRCC oncogenesis, which may contribute to available knowledge, improve the treatment designs, and enhance the accuracy of prognosis for patients with ccRCC.

## 7 Author Contributions

NJ and YLZ developed this idea and designed this research. NJ, YLZ, ZHZ and THL analysed the data. NJ, YLZ, ZHZ and THL wrote the draft of the manuscript. HQG and RY obtained copies of studies and revised the writing. All authors have read and agreed to the published version of the manuscript.

## 9 Acknowledgments

None.

## 10 Data Availability Statement

The datasets for this study can be found in the online databases.

## Reference

1. Capitanio U, Bensalah K, Bex A, et al. Epidemiology of Renal Cell Carcinoma. Eur Urol. Jan 2019;75(1):74–84. doi:10.1016/j.eururo.2018.08.036

2. Ferlay J, Soerjomataram I, Dikshit R, et al. Cancer incidence and mortality worldwide: sources, methods and major patterns in GLOBOCAN 2012. Int J Cancer. Mar 1 2015;136(5):E359–86. doi:10.1002/ijc.29210

3. Capitanio U, Montorsi F. Renal cancer. Lancet. Feb 27 2016;387(10021):894–906. doi:10.1016/S0140-6736(15)00046-X

4. Pierorazio PM, Johnson MH, Patel HD, et al. Management of Renal Masses and Localized Renal Cancer: Systematic Review and Meta-Analysis. J Urol. Oct 2016;196(4):989–99. doi:10.1016/j.juro.2016.04.081

5. Sun Y, Ponz-Sarvise M, Chang SS, et al. Proteasome inhibition enhances the killing effect of BikDD gene therapy. Am J Transl Res. 2015;7(2):319–27.

6. Wang CY, Li CY, Hsu HP, et al. PSMB5 plays a dual role in cancer development and immunosuppression. Am J Cancer Res. 2017;7(11):2103–2120.

7. O’Hurley G, O’Grady A, Smyth P, et al. Evaluation of Zinc-alpha-2-Glycoprotein and Proteasome Subunit beta-Type 6 Expression in Prostate Cancer Using Tissue Microarray Technology. Appl Immunohistochem Mol Morphol. Dec 2010; 18(6):512–7. doi:10.1097/PAI.0b013e3181e29998

8. Sorokin AV, Kim ER, Ovchinnikov LP. Proteasome system of protein degradation and processing. Biochemistry (Mosc). Dec 2009;74(13):1411–42. doi:10.1134/s000629790913001x

9. Yuan F, Ma Y, You P, et al. A novel role of proteasomal beta1 subunit in tumorigenesis. BiosciRep. Jul 16 2013;33(4)doi:10.1042/BSR20130013

10. Yang G, Jian L, Lin X, Zhu A, Wen G. Bioinformatics Analysis of Potential Key Genes in Trastuzumab-Resistant Gastric Cancer. Dis Markers. 2019;2019:1372571. doi:10.1155/2019/1372571

11. Wang M, Windgassen D, Papoutsakis ET. Comparative analysis of transcriptional profiling of CD3+, CD4+ and CD8+ T cells identifies novel immune response players in T-cell activation. BMC Genomics. May 16 2008;9:225. doi:10.1186/1471-2164-9-225

12. Chandrashekar DS, Bashel B, Balasubramanya SAH, et al. UALCAN: A Portal for Facilitating Tumor Subgroup Gene Expression and Survival Analyses. Neoplasia. Aug 2017;19(8):649–658. doi:10.1016/j.neo.2017.05.002

13. Asplund A, Edqvist PH, Schwenk JM, Ponten F. Antibodies for profiling the human proteome-The Human Protein Atlas as a resource for cancer research. Proteomics. Jul 2012;12(13):2067–77. doi:10.1002/pmic.201100504

14. Gyorffy B, Surowiak P, Budczies J, Lanczky A. Online survival analysis software to assess the prognostic value of biomarkers using transcriptomic data in non-small-cell lung cancer. PLoS One. 2013;8(12):e82241. doi:10.1371/journal.pone.0082241

15. Gao J, Aksoy BA, Dogrusoz U, et al. Integrative analysis of complex cancer genomics and clinical profiles using the cBioPortal. Sci Signal. Apr 2 2013;6(269):pl1. doi:10.1126/scisignal.2004088

16. Zhou Y, Zhou B, Pache L, et al. Metascape provides a biologist-oriented resource for the analysis of systems-level datasets. Nat Commun. Apr 3 2019;10(1):1523. doi:10.1038/s41467-019-09234-6

17. Franz M, Rodriguez H, Lopes C, et al. GeneMANIA update 2018. Nucleic Acids Res. Jul 2 2018;46(W1):W60–W64. doi:10.1093/nar/gky311

18. Szklarczyk D, Gable AL, Lyon D, et al. STRING v11: protein-protein association networks with increased coverage, supporting functional discovery in genome-wide experimental datasets. Nucleic Acids Res. Jan 8 2019;47(D1):D607–D613. doi:10.1093/nar/gky1131

19. Huang da W, Sherman BT, Lempicki RA. Systematic and integrative analysis of large gene lists using DAVID bioinformatics resources. Nat Protoc. 2009;4(1):44–57. doi:10.1038/nprot.2008.211

20. Li T, Fan J, Wang B, et al. TIMER: A Web Server for Comprehensive Analysis of Tumor-Infiltrating Immune Cells. Cancer Res. Nov 1 2017;77(21):e108–e110. doi:10.1158/0008-5472.CAN-17-0307

21. Chen L, Madura K. Increased proteasome activity, ubiquitin-conjugating enzymes, and eEF1A translation factor detected in breast cancer tissue. Cancer Res. Jul 1 2005;65(13):5599–606. doi:10.1158/0008-5472.CAN-05-0201

22. Varga G, Mikala G, Kiss KP, et al. Proteasome Subunit Beta Type 1 P11A Polymorphism Is a New Prognostic Marker in Multiple Myeloma. Clin Lymphoma Myeloma Leuk. Nov 2017;17(11):734–742. doi:10.1016/j.clml.2017.06.034

23. Walter RFH, Sydow SR, Berg E, et al. Bortezomib sensitivity is tissue dependent and high expression of the 20S proteasome precludes good response in malignant pleural mesothelioma. Cancer Manag Res. 2019;11:8711–8720. doi:10.2147/CMAR.S194337

24. Wu F, Niu Z, Zhou B, Li P, Qian F. PSMB1 Negatively Regulates the Innate Antiviral Immunity by Facilitating Degradation of IKK-epsilon. Viruses. Jan 24 2019;11(2) doi:10.3390/v11020099

25. Tan S, Li H, Zhang W, et al. NUDT21 negatively regulates PSMB2 and CXXC5 by alternative polyadenylation and contributes to hepatocellular carcinoma suppression. Oncogene. Aug 2018;37(35):4887–4900. doi:10.1038/s41388-018-0280-6

26. Dumartin L, Whiteman HJ, Weeks ME, et al. AGR2 is a novel surface antigen that promotes the dissemination of pancreatic cancer cells through regulation of cathepsins B and D. Cancer Res. Nov 15 2011;71(22):7091–102. doi:10.1158/0008-5472.CAN-11-1367

27. Weyburne ES, Wilkins OM, Sha Z, et al. Inhibition of the Proteasome beta2 Site Sensitizes Triple-Negative Breast Cancer Cells to beta5 Inhibitors and Suppresses Nrf1 Activation. Cell Chem Biol. Feb 16 2017;24(2):218–230. doi:10.1016/j.chembiol.2016.12.016

28. Gemoll T, Habermann JK, Becker S, et al. Chromosomal aneuploidy affects the global proteome equilibrium of colorectal cancer cells. Anal Cell Pathol (Amst). 2013;36(5-6):149–61. doi:10.3233/ACP-140088

29. Wang H, He Z, Xia L, et al. PSMB4 overexpression enhances the cell growth and viability of breast cancer cells leading to a poor prognosis. Oncol Rep. Oct 2018;40(4):2343–2352. doi:10.3892/or.2018.6588

30. Guo S, Chen Y, Yang Y, et al. TRIB2 modulates proteasome function to reduce ubiquitin stability and protect liver cancer cells against oxidative stress. Cell Death Dis. Jan 7 2021;12(1):42. doi:10.1038/s41419-020-03299-8

31. Munkacsy G, Abdul-Ghani R, Mihaly Z, et al. PSMB7 is associated with anthracycline resistance and is a prognostic biomarker in breast cancer. Br J Cancer. Jan 19 2010;102(2):361–8. doi:10.1038/sj.bjc.6605478

32. Cao Y, Tang W, Tang W. Immune cell infiltration characteristics and related core genes in lupus nephritis: results from bioinformatic analysis. BMC Immunol. Oct 21 2019;20(1):37. doi:10.1186/s12865-019-0316-x

33. Nishida K, Kawashima A, Kanazawa T, et al. Clinical importance of the expression of CD4+CD8+ T cells in renal cell carcinoma. Int Immunol. May 8 2020;32(5):347–357. doi:10.1093/intimm/dxaa004

34. Sarrabay G, Mechin D, Salhi A, et al. PSMB10, the last immunoproteasome gene missing for PRAAS. J Allergy Clin Immunol. Mar 2020;145(3):1015–1017 e6. doi:10.1016/j.jaci.2019.11.024

